# Regulation of cancer epigenomes with a histone-binding synthetic transcription factor

**DOI:** 10.1101/072975

**Authors:** David B. Nyer, Daniel Vargas, Caroline Hom, Karmella A. Haynes

## Abstract

Chromatin proteins have expanded the mammalian synthetic biology toolbox by enabling control of active and silenced states at endogenous genes. Others have reported synthetic proteins that bind DNA and regulate genes by altering chromatin marks, such as histone modifications. Previously we reported the first synthetic transcriptional activator, the "Polycomb-based transcription factor" (PcTF), that reads histone modifications through a protein-protein interaction between the PCD motif and trimethylated lysine 27 of histone H3 (H3K27me3). Here, we describe the genome-wide behavior of PcTF. Transcriptome and chromatin profiling revealed PcTF-sensitive promoter regions marked by proximal PcTF and distal H3K27me3 binding. These results illuminate a mechanism in which PcTF interactions bridge epigenetic marks with the transcription initiation complex. In three cancer-derived human cell lines tested here, many PcTF-sensitive genes encode developmental regulators and tumor suppressors. Thus, PcTF represents a powerful new fusion-protein-based method for cancer research and treatment where silencing marks are translated into direct gene activation.

## Introduction

Proteins from the gene regulatory complex known as chromatin mediate stable, epigenetic expression states that persist over multiple cell divisions in metazoan tissues. Harnessing the potent gene-regulating functions of chromatin proteins has become a high-priority for cancer therapy and tissue engineering. The “histone code” model of chromatin function^1, 2^ has strongly influenced work in epigenetic engineering and drug development^3, 4^. According to this model, biochemical marks are written onto DNA-bound histone proteins and these marks are read when effector proteins physically interact with the modified histones. Then the effector proteins enhance or inhibit transcription initiation. Much of our current understanding about chromatin-mediated gene regulation comes from deconstructive methods such as genetic mutations and RNA interference (for examples see^5, 6, 7^). Constructive approaches, where synthetic systems are built from chromatin components, are gaining recognition as an important and powerful research method^8, 9^ as well as a powerful application for biomedical engineering.

We constructed a synthetic transcription factor based on the natural effector protein CBX8. In many human cancers, lysine 27 on histone H3 (H3K27) becomes marked by trimethylation at abnormally high levels near tumor suppressor loci. CBX8-containing complexes accumulate at H3K27me3 and repress gene transcription^10, 11^. A central step in the Polycomb pathway is the specific interaction between the H3K27me3 mark and a hydrophobic binding pocket within the Polycomb chromodomain (PCD) motif of CBX isoforms^12, 13^. We used this interaction to design a Polycomb-based transcription factor (PcTF) that has VP64 (tetrameric VP16) and a visible red fluorescent tag (mCherry) fused to the C-terminus of the 60 amino acid PCD. The C-terminal VP64 domain allows PcTF to stimulate activation at repressed H3K27me3-associated genes (Fig. 1a). In previous work, we demonstrated H3K27me3-dependent PcTF activity at a model locus (*UASTk-luciferase*). PcTF also activated endogenous silenced loci in an osteosarcoma cell line (U-2 OS)^14^. Low-resolution protein mapping experiments at four loci identified two promoter regions that were co-occupied by PcTF and H3K27me3. Further investigation is needed to understand and accurately predict targets of the PcTF transcription activator. Here, we report genome-wide analyses that substantially advance our understanding of PcTF function. PcTF-stimulated gene activation in three different cancer cell types identified a large cohort of universally upregulated genes with a signature H3K27me3 enrichment profile. PcTF is enriched at transcription start sites within the nucleosome-free region of promoters and this profile depends upon the methyl-histone-binding PCD domain. We present a model where PcTF bridges distal H3K27me3 with endogenous transcription factors near the transcription start site. These findings provide significant progress towards predicting targets of PcTF based on distributions of epigenetic marks.

**Figure 1.**
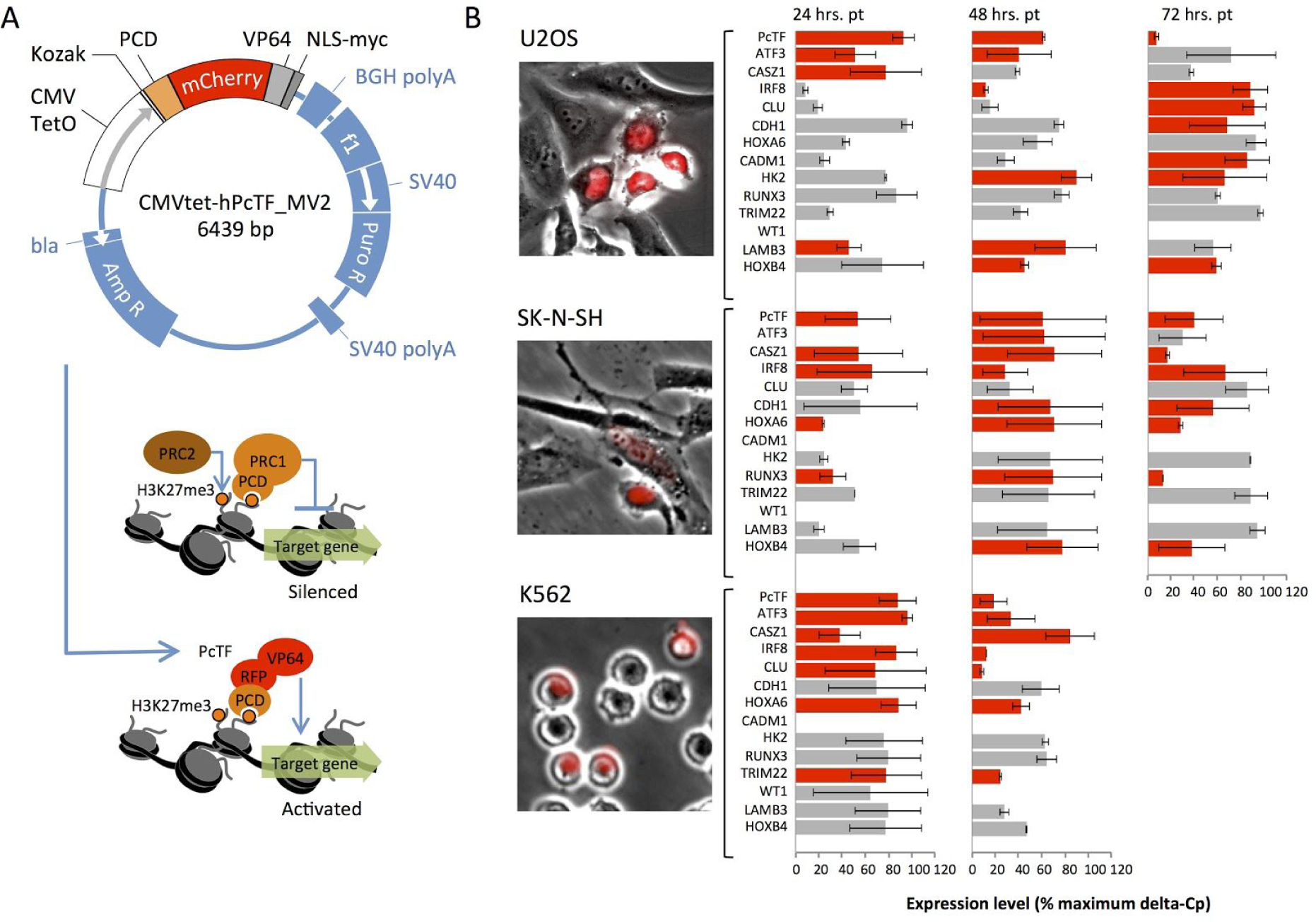
PcTF expression stimulates upregulation of known targets of Polycomb in transiently transfected cells. (A) Map of the PcTF-expressing plasmid (top). The natural PRC1 complex mediates gene silencing (middle). PcTF expression leads to accumulation of PcTF at H3K27me3 and gene activation (bottom)^14^. (B) Transiently-transfected, PcTF-positive U-2 OS, SK-N-SH and K562 cells are visualized via the red fluorescent protein tag mCherry (RFP). qRT-PCR was used to determine mRNA levels of PcTF and a panel of 13 target genes. Red bars represent 2-fold or greater GAPDH-normalized expression in PcTF-plasmid transfected cells compared to cells mock-transfected with the vehicle (Lipofectamine LTX) only. Hrs. pt = hours post-transfection. Error bars = standard deviation for two independent transfections.

## RESULTS

### Common Polycomb-silenced loci are activated in PcTF-expressing cells

We measured PcTF-mediated activation of known Polycomb targets in three different cell lines, osteosarcoma (U-2 OS), neuroblastoma (SK-N-SH), and leukemia (K562). We selected a panel of 13 genes for which Polycomb-mediated silencing is supported by genetic and pharmacologic disruption studies^15, 16, 17, 18, 19, 20, 21, 22, 23, 24, 25^ and protein mapping studies^26, 27, 28, 29^ in human cancers and stem cells: *ATF3*, *CADM1*, *CASZ1*, *CDH1*, *CLU*, *HK2*, *HOXA6*, *HOXB4*, *IRF8*, *LAMB3*, *RUNX3*, *TRIM22*, and *WT1*. We transfected U2-OS, SK-N-SH, and K562 cells with a PcTF-expressing plasmid (Fig. 1a) via Lipofectamine LTX. We used quantitative reverse transcription PCR (qRT-PCR) to measure mRNA from mock-transfected (vehicle only) and plasmid-transfected cells (Fig. 1b). Primers and dye-conjugated hydrolysis probes are described in Table S1. In all three cell lines, over half of the genes were activated two-fold or higher compared to mock-transfected controls. Up-regulation persisted for several genes as PcTF transcript levels decreased over time. Three of the Polycomb target genes were upregulated in all three cell types: *ATF3*, *CASZ1*, and *IRF8*.

We used data from chromatin immunoprecipitation followed by deep sequencing (ChIP-seq) to detect H3K27me3 enrichment at Polycomb target genes. In U-2 OS, ChIP was performed as previously described using an antibody against H3K27me3^14^. DNA was purified from the immunoprecipitated chromatin, analyzed by next generation deep sequencing, and reads were aligned to the hg19 human genome consensus (Feb. 2009 GRCh37). Regions of enrichment compared to non-immunoprecipitated bulk chromatin (input) were calculated using the Hotspot algorithm^30^. We used shared and public data to determine H3K27me3 enrichment in SK-N-SH and K562 cells. *ATF3*, *CASZ1*, and *IRF8* are marked by H3K27me3 (Fig. S1) and become upregulated upon PcTF expression in all three cell types tested here. *HOXA6*, *HK2*, and *TRIM22* are enriched for H3K27me3 in one or two cell types, and only become up-regulated when H3K27me3 is present (Fig. S1). These results are consistent with a mechanism where PcTF accumulates at H3K27me3 sites and induces gene transcription.

### PcTF stimulates expression of a large subset of H3K27me3-marked genes in U-2 OS, SK-N-SH, and K562 cells

Whole-transcriptome analysis confirmed up-regulation of the known Polycomb targets and revealed a larger subset of commonly up-regulated genes. We performed next generation deep sequencing of total RNA (RNA-seq) from mock transfected (control) and PcTF-expressing cells (+PcTF) up to 96 hours post-transfection. Whole-transcriptome analyses of U-2 OS, SK-N-SH, and K562 corroborated the qRT-PCR results for 9 of the 13 known Polycomb targets: *ATF3*, *CASZ1*, *CLU*, *HK2*, *HOXA6*, *HOXB4*, *IRF8*, *LAMB3*, and *TRIM22*. Comparison of fragments per kilobase of gene model per million mapped reads (FPKM) for control versus +PcTF samples showed that 10.3 - 23.9% of 23,245 annotated Refseq genes became up-regulated 2-fold or higher (Fig. 2a). This result is consistent with a mechanism where PcTF accumulates at silenced, H3K27me3-enriched loci and activates transcription through its C-terminal VP64 domain. We also observed down-regulation (fold-change ≤ 2.0) at 6.2 - 9.3% of the genes in the three cell lines we tested. Repression is inconsistent with the transcriptional activation function of VP64. In some cases, PcTF might disrupt proper gene expression by placing VP64 in a suboptimal position relative to the promoter^31, 32^.

**Figure 2.**
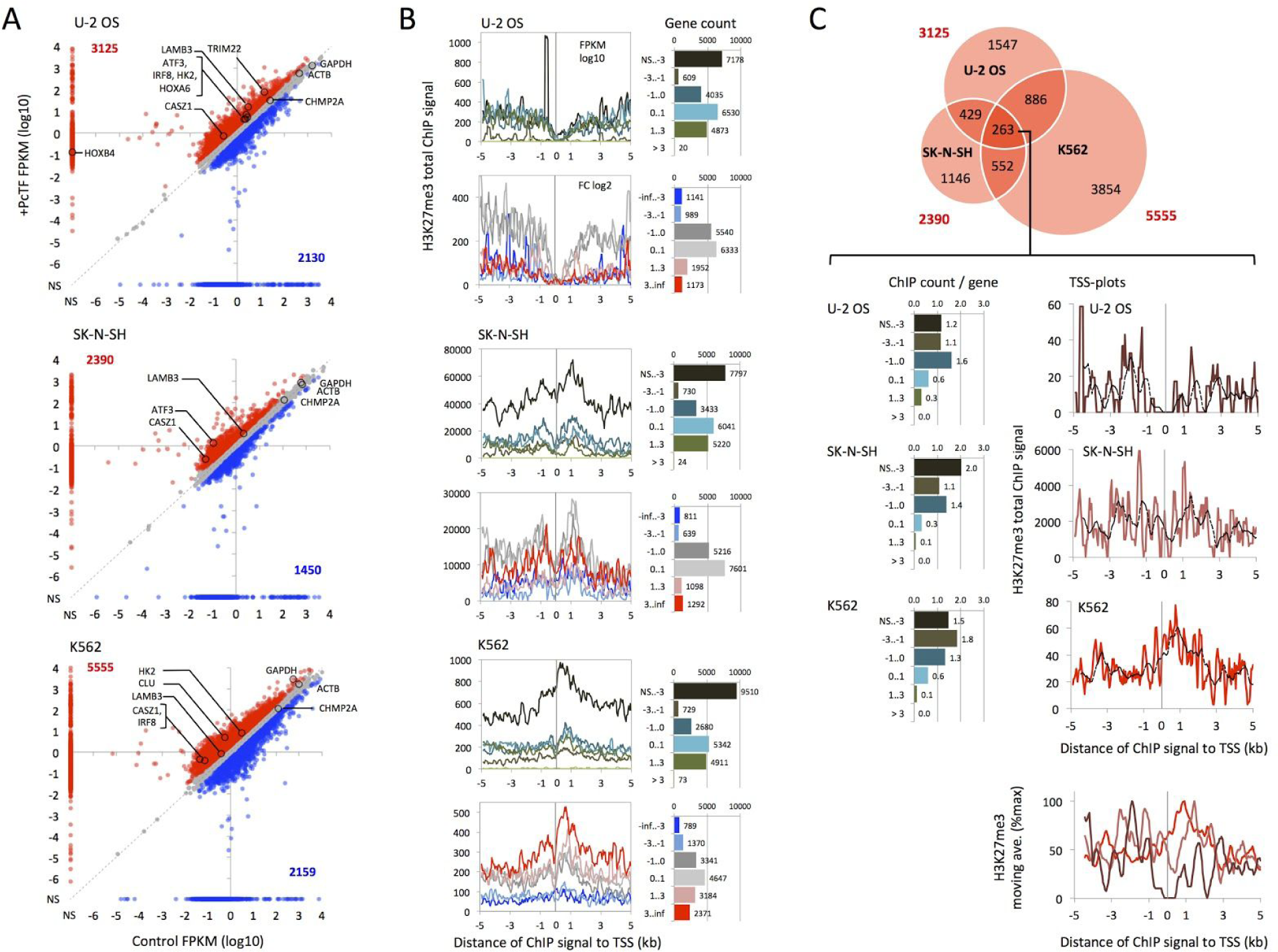
Genome-wide analysis of gene transcription and H3K27me3 at promoter regions in PcTF-expressing cells. (A) Scatter plots compare RNA-seq signals (FPKM log10) of 23,245 genes from cells that were mock transfected (Control) or expressed PcTF (+PcTF) 96 hours post-transfection for U2-OS and SK-N-SH and 48 hours for K562. Active housekeeping genes GAPDH, ACTB, and CHMP2A are negative controls. NS = no signal. (B) TSS plots show total H3K27me3 ChIP enrichment values mapped at 200 bp intervals with a step value of 50 bp. Values are stratified by gene expression level (FPKM log10) in the untreated (no PcTF) state or by fold change (FC log2) after PcTF treatment and normalized by the frequency of genes in each category. Genes where FPKM = 0 for both control and treated cells were excluded from the FC log2-stratified data. (C) The Venn diagram shows unique and common sets of genes that become up-regulated at least two-fold in the three cell types. Moving averages for each TSS plot (dashed line, window = 10) are scaled and overlaid in the bottom plot.

We looked for an epigenetic signature of PcTF-sensitive genes by mapping enrichment of H3K27me3 near transcription start sites (TSS’s). Overall, H3K27me3 enrichment was depleted near transcription start sites at silenced and active genes, indicated by a sharp valley at position zero in stratified TSS-plots (Fig. 2b). This profile has been observed in other reports; similar plots show a valley in the TSS profile at the nucleosome-free region (NFR) of promoters in mammalian cells^33, 34^. Genes with lower basal expression levels, prior to PcTF expression, show the highest levels of H3K27me3 enrichment which is consistent with the role of H3K27me3 in gene silencing. Genes that became up-regulated in response to PcTF expression showed the highest H3K27me3 levels compared to non-responsive and down-regulated genes in K562 cells. This result provides evidence of a mechanism where PcTF accumulates at methyl-histone marks through the N-terminal PCD and activates silenced genes through the C-terminal VP64 domain. In SK-N-SH and U-2 OS however, up-regulated genes showed the same or lower H3K27me3 levels as non-responsive genes. Therefore, PcTF-sensitivity may not be strictly determined by methylation signal levels that were determined by ChIP analysis.

We hypothesized that a specific H3K27me3 distribution pattern around the TSS determines which genes are activated by PcTF. We identified a subset of 263 commonly upregulated genes (Fig. 2c) in order to exclude genes that behave in a cell-type-specific manner. In U-2 OS and SK-N-SH, we observed depletion of H3K27me3 at the TSS, surrounded by signal peaks at intervals of roughly 1 kb. ChIP hotspot densities around the TSS were highest for genes with low basal expression levels (0.001 - 1.0 FPKM) (Fig. 2c). The highest peaks for K562 were observed immediately downstream of the TSS and 3 kb upstream (Fig. 2c). Altogether, these data suggest that PcTF-mediated activation of silenced genes may rely on H3K27me3 enrichment at positions distal to the TSS.

### ChIP-seq analyses show PCD-dependent enrichment of PcTF near transcription start sites

In order to further investigate the mechanism of PcTF-mediated gene regulation, we used ChIP-seq to compare the distribution of PcTF and H3K27me3 throughout the genome and at TSS regions. We performed ChIP-seq on chromatin from an isogenic PcTF-expressing U-2 OS cell line. The cell line “U2OS-PcTF” carries a chromosomally-integrated, TetR-repressed gene that expresses myc-tagged PcTF when the cells are treated with doxycycline (Fig. 3a), as described previously^14^. We used this system for ChIP-seq instead of transiently transfected cells in order to eliminate negative background from PcTF-minus cells. ChIP was performed as previously described using either an antibody against the PcTF N-terminal myc tag for U2OS-PcTF cells or anti-H3K27me3 for the Flp-in T-REx parental line^14^. ChIP-seq analysis of a truncated version of PcTF that lacks the PCD domain (ΔTF) showed little overlap with PcTF and H3K27me3 (Fig. S2), which indicated that PcTF and H3K27me3 distributions were not due to random crosslinking. We analyzed the distributions of PcTF and H3K27me3 ChIP at different genomic features and observed the greatest enrichment over background for intergenic regions (Fig. 3b). PcTF, but not H3K27me3, showed signal enrichment at promoters. These results indicate that PcTF and H3K27me3 co-occupy non-coding regions, and that PcTF is frequently enriched at promoter sites.

**Figure 3.**
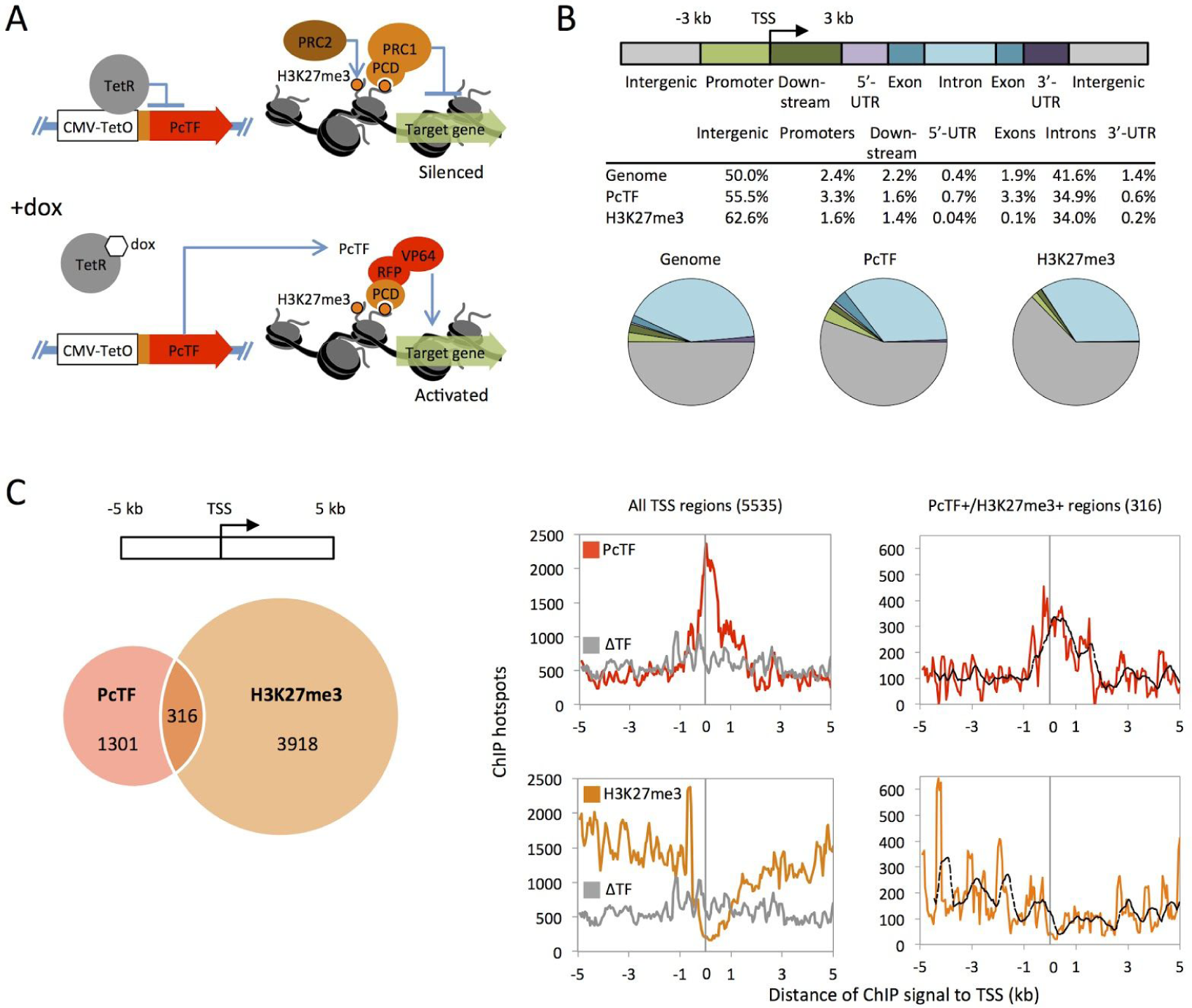
ChIP-seq analysis of PcTF and H3K27me3 distribution in a transgenic U2-OS cell line. (A) Illustrated mechanism of PcTF function. Doxycycline (dox) induced PcTF expression in U2OS-PcTF cells leads to accumulation of PcTF at H3K27me3-positive promoters and gene activation^14^. (B) Distribution of PcTF and H3K27me3 ChIP signals (hotspots) in genes and intergenic regions compared to background (Genome) distributions. (C) The venn diagram compares genes that have PcTF- and H3K27me3-marked TSS regions. TSS-centered plots (left) show the total ChIP hotspot signal value within a sliding window of 200 bp (step size = 50 bp). ΔTF is a negative control protein that lacks the PCD (H3K27me3-binding) motif. The right-most plots show the subset of 316 genes where the TSS region is co-occupied by PcTF and H3K27me3. Dashed lines show the moving average (window = 10).

We inspected promoter regions more closely by identifying PcTF and H3K27me3 enriched regions surrounding TSS’s. We identified 5535 regions that were marked with PcTF or H3K27me3 or both. A TSS plot showed a peak of PcTF enrichment at the TSS over a 2 kb interval (Fig. 3c). Randomly crosslinked ΔTF did not show the same enrichment profile, therefore mCherry-tagged VP64 alone does not account for the enrichment pattern of PcTF. In contrast to PcTF, H3K27me3 was depleted at the TSS. The subset of 316 regions in which we detected both PcTF and H3K27me3 showed a similar but shorter PcTF enrichment peak. H3K27me3 signal was depleted near the TSS, with dispersed peaks of H3K27me3 further from the TSS. The profiling data suggest that at co-occupied regions, PcTF enrichment appears at and adjacent to the TSS, while H3K27me3 is farther away either upstream or downstream. Overall, these results suggest that PcTF accumulates near the TSS, and that this accumulation depends upon interaction with distal H3K27me3 through the N-terminal PCD peptide.

### Up-regulated genes are marked by H3K27me3 upstream and PcTF near the TSS in U2OS-PcTF cells

Up-regulation of PcTF-sensitive genes corresponds with H3K27me3 enrichment at roughly 1 - 2 kb upstream of the transcription start site in regions where we also detected PcTF enrichment near the TSS. To identify up-regulated genes, we compared RNA-seq FPKM values of 23,244 genes for U2OS-PcTF cells that were untreated or treated with doxycycline (dox) to induce PcTF expression. Known Polycomb targets *LAMB3*, *IRF8*, *CASZ1*, *RUNX3*, *HOXB4*, and *CDH1* were up-regulated 2-fold or higher. For transcript levels that changed at least two-fold in either direction, the predominant response was an increase in expression of genes that had a low initial expression state in control cells. FPKM values of silenced genes are typically lower than 1.0^35^. The frequency of upregulated genes within the non-expressing FPKM range of 10^-4^ - 10^0^ was 23%, whereas only 3% of active genes were upregulated (Fig. 4a).

**Figure 4.**
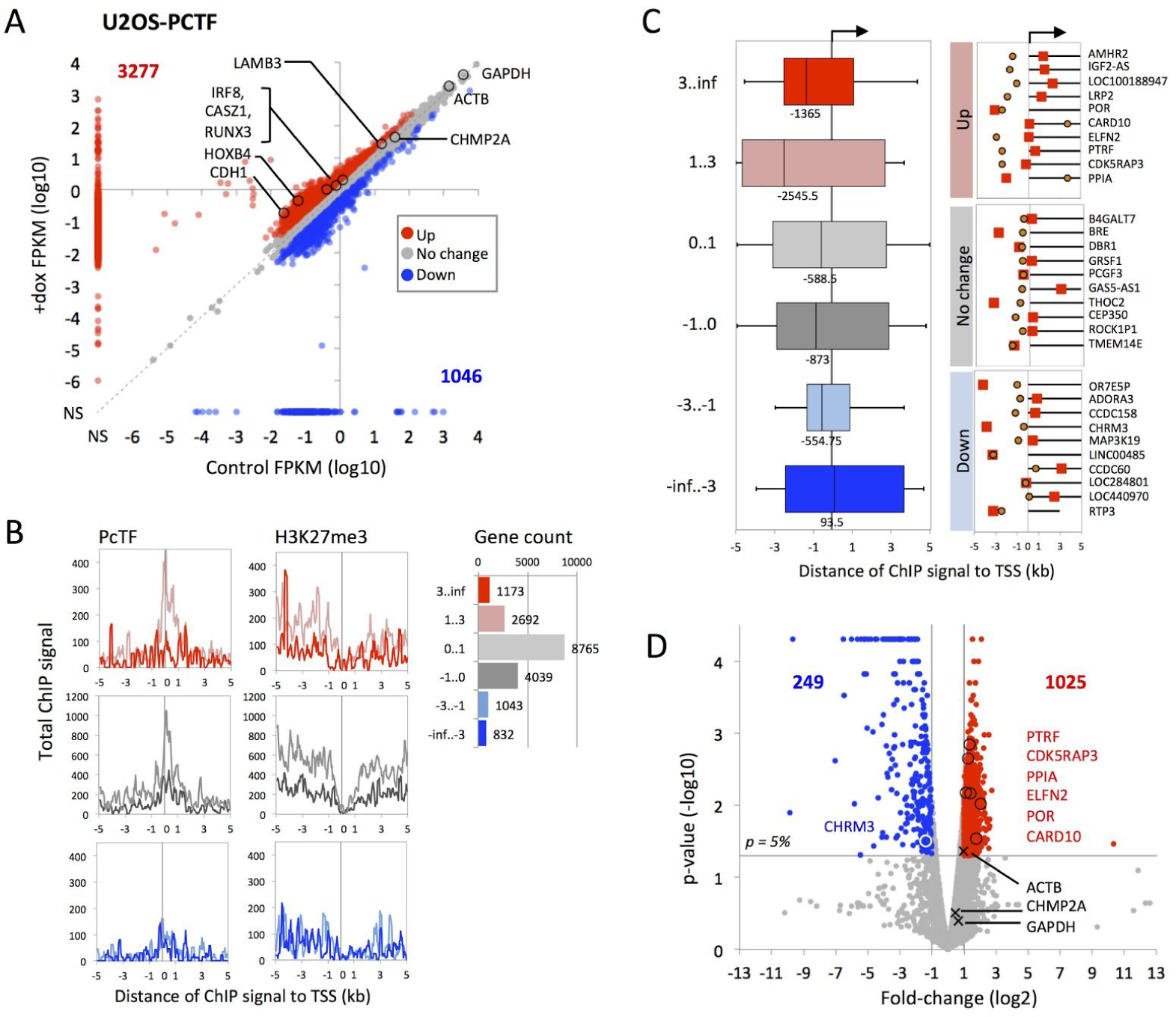
Analysis of PcTF and H3K27me3 localization and gene regulation. (A) The x/y scatter plot compares RNA-seq signals (FPKM log10) of U2OS-PcTF cells without PcTF (Control) and after PcTF induction (+dox). Known Polycomb-regulated genes and control genes from Figures 1 and 2 are highlighted for comparison with previous experiments. (B) TSS-centered plots show the total ChIP hotspot signal value (window = 200 bp, step size = 50 bp) stratified by log2 fold-change expression. (C) H3K27me3 ChIP signal distances from the TSS were compared for genes of lengths ≥ 2 kb in the subset of 316 genes where the 10 kb region is co-occupied by PcTF and H3K27me3. Genes were grouped by fold-change expression (as in B). Box plots show median values (solid vertical line), 25th (left box) and 75th (right box) percentiles, and minimum (left whisker) and maximum values (right whisker). TSS maps drawn to scale show the midpoints of ChIP signals for H3K27me3 (orange circle) and PcTF (red square) relative to the TSS. Genes are sorted from highest to lowest log2 fold-change value. (D) The volcano plot shows statistical significance versus fold change (same color scheme as in Fig. 1a) for duplicate RNA-seq experiments. Up- and down-regulated genes from panel C are marked with circles. Control genes are marked with an “x.”

Stratified TSS plots showed that PcTF was frequently enriched immediately downstream of the TSS at up-regulated genes (Fig. 4b). H3K27me3 was depleted near the TSS and showed peaks of enrichment 1 kb, 2 kb and 4 kp upstream. Median distances of the nearest H3K27me3 signal were -2545.5 bp for 2- to 8-fold up-regulated genes and -1365 bp for genes that were up-regulated 8-fold or higher (Fig. 4c). We observed a significant (p < 0.05) increase in expression across two independent experimental replicates for five of the upregulated genes: *PTRF*, *CDKRAP3*, *PPIA*, *ELFN2*, *POR*, and *CARD10* (Fig. 4d). For genes that showed no change or were down-regulated, median distances ranged from -588.5 to 93.5 bp. Positioning of H3K27me3 very close to the TSS at non-responsive and down-regulated genes might interfere with VP64-driven gene activation, or cause VP64 to recruit transcription factors to a site that is not optimal for proper transcription. These data indicate that a distal H3K27me3 mark positions PcTF at an optimal position to allow the VP64 activation domain to stimulate gene transcription at target promoters.

### H3K27me3 enrichment occurs near enhancers at PcTF-sensitive loci

Sites that are distal from the TSS are known as enhancers, which are often bound by regulatory proteins that interact with transcription factors at the TSS and stimulate or repress gene expression. We hypothesized that PcTF stimulates expression at genes where a nearby enhancer is enriched for H3K27me3. Investigation of the chromosomal positions of PcTF-sensitive genes, enhancers, and H3K27me3 revealed that histone methylation at upstream enhancers corresponds with PcTF-mediated gene activation. Enhancer regions were identified from the 15 classes of human chromatin states from the ENCODE project for 9 different human cell types^36^, including K562. Enhancer coordinates were based on class 4 (strong enhancer) and class 5 (medium enhancer) hg19 annotations from the public K562 cell data. We identified the closest enhancer element near 23,245 TSS’s at both the upstream and downstream position and calculated median distances to the nearest H3K27me3 signal (Fig. 5a). Genes with FPKM values of zero in both control and treated cells were excluded from the calculations. We observed that H3K27me3 ChIP signals occur close to upstream enhancer midpoints at higher frequencies near up-regulated genes. U-2 OS cells showed the strongest correlation (Pearson R^2^ = 0.79) between H3K27me3-proximal, upstream, class 4 enhancers and fold-change expression in response to PcTF.

**Figure 5.**
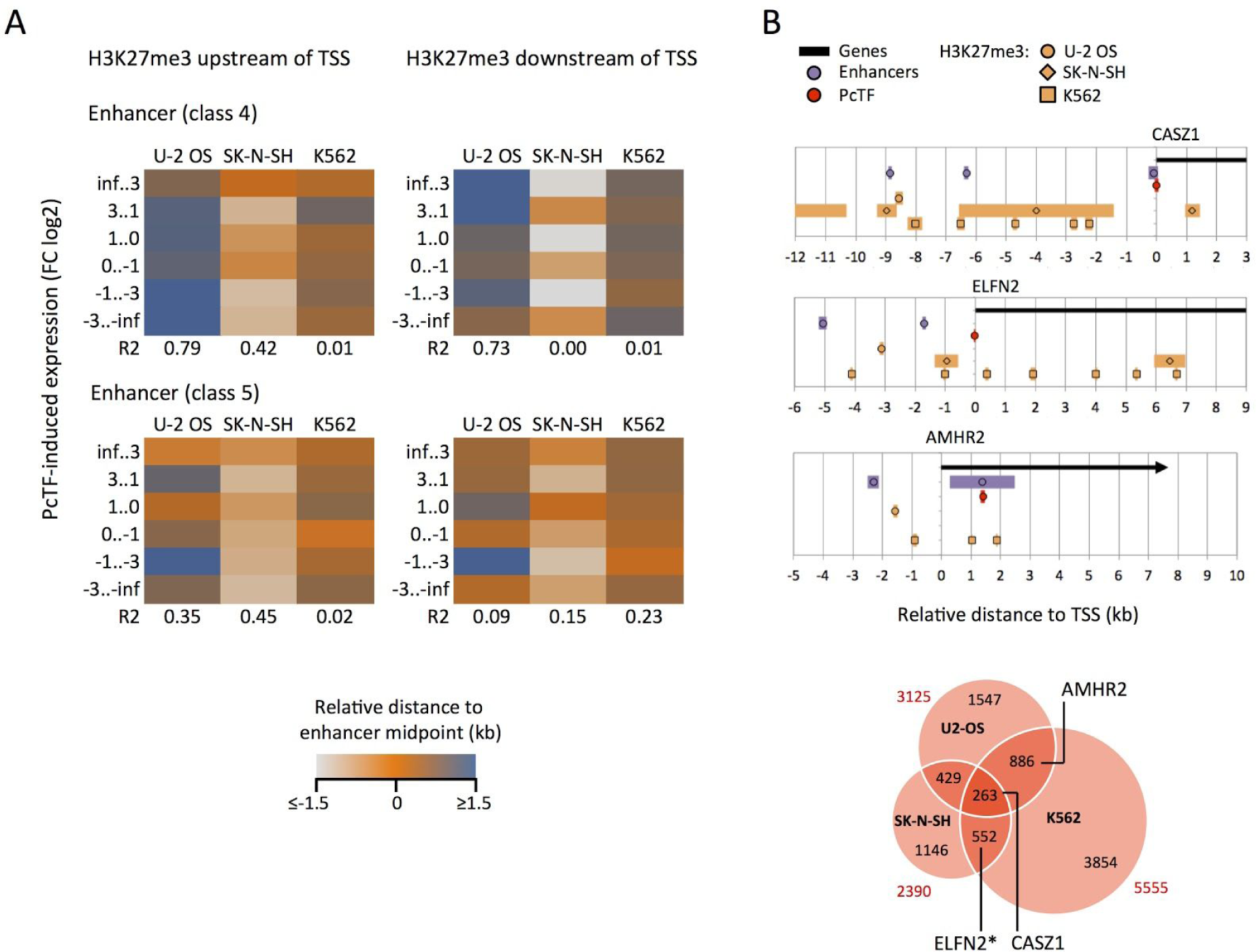
H3K27me3 enrichment at enhancer regions. (A) Color scales show median distances between midpoints of nearest H3K27me3 ChIP hotspots and enhancer elements closest to the TSS of all genes. Data are stratified by fold-change in expression level after PcTF expression. Enhancer coordinates were determined using chromatin states classes from the ENCODE project (UCSC Browser HMM track for K562 cells)^36^. R2 = Pearson correlation coefficient of median distance versus fold-change bin. (B) High-resolution maps of features at select genes that are upregulated in one or more cell lines. The Venn diagram from Fig. 2c is shown here with labels for genes that are shown in the maps. *ELFN was up-regulated in U2OS-PcTF and not in transiently transfected U-2 OS cells.

Close inspection of gene regions further supported the significance of H3K27me3 positioning at enhancers near PcTF-sensitive genes. At *CASZ1*, one of the known Polycomb targets, the upstream enhancer regions were positioned near H3K27me3 enrichment signals in all three cell types. *ELFN2*, a gene that became up-regulated after PcTF expression in all three cell types, also showed H3K27me3 proximity to upstream enhancers. At *AMHR2*, H3K27me3 ChIP hotspots appear about 1 kb from the TSS, between the nearest enhancer and the nearest PcTF ChIP signal in U-2 OS and K562 (Fig. 5b). *AMHR2* was up-regulated by PcTF in U-2 OS and K562, but not in SK-N-SH where we found no H3K27me3 signal within the same region. These results suggest that H3K27me3 at enhancer elements upstream of the TSS contributes to PcTF-mediated gene activation.

## DISCUSSION

The work presented here provides new insights into the mechanism of PcTF, a synthetic chromatin-based transcription factor. PcTF couples recognition of a silencing-associated mark with gene activation. PcTF enrichment near TSS sites is similar to the patterns that have been observed for subunits of the transcription initiation complex^33, 37^, suggesting a strong interaction with endogenous transcription factors. The C-terminal domain of PcTF includes VP64 which is composed of four copies of a core acidic transcription activation domain (TAD) from VP16^38^. The TAD is derived from the H1 region of VP16, which has been shown to bind with high affinity (high nanomolar range) to the Mediator complex subunit MED25^39^. The VP16 TAD also interacts with MED17, several members of the Transcription Factor II (TFII) family, and the TATA promoter motif-binding protein TBP (reviewed in^40, 41^). In contrast, the C-terminal PCD has relatively low affinity (high micromolar range) for H3K27me3^42^. PCD-H3K27me3 binding may be weak *in vivo*, but it is still necessary for PcTF function^14^ and for TSS-proximal enrichment (Fig. 3C). Enrichment of PcTF in regions that are near to but do not overlap with H3K27me3 was observed in our previous ChIP-PCR studies^14^. This offset of PcTF ChIP signals from H3K27me3 sites suggests that PcTF engages with H3K27me3 and then becomes trapped near the TSS through interactions with the transcription initiation proteins in crosslinked chromatin. The data provide strong evidence for a mechanism where PcTF bridges distal histone methylation marks with PolII-associated transcription factors at the nucleosome-free TSS (Fig. 6).

**Figure 6.**
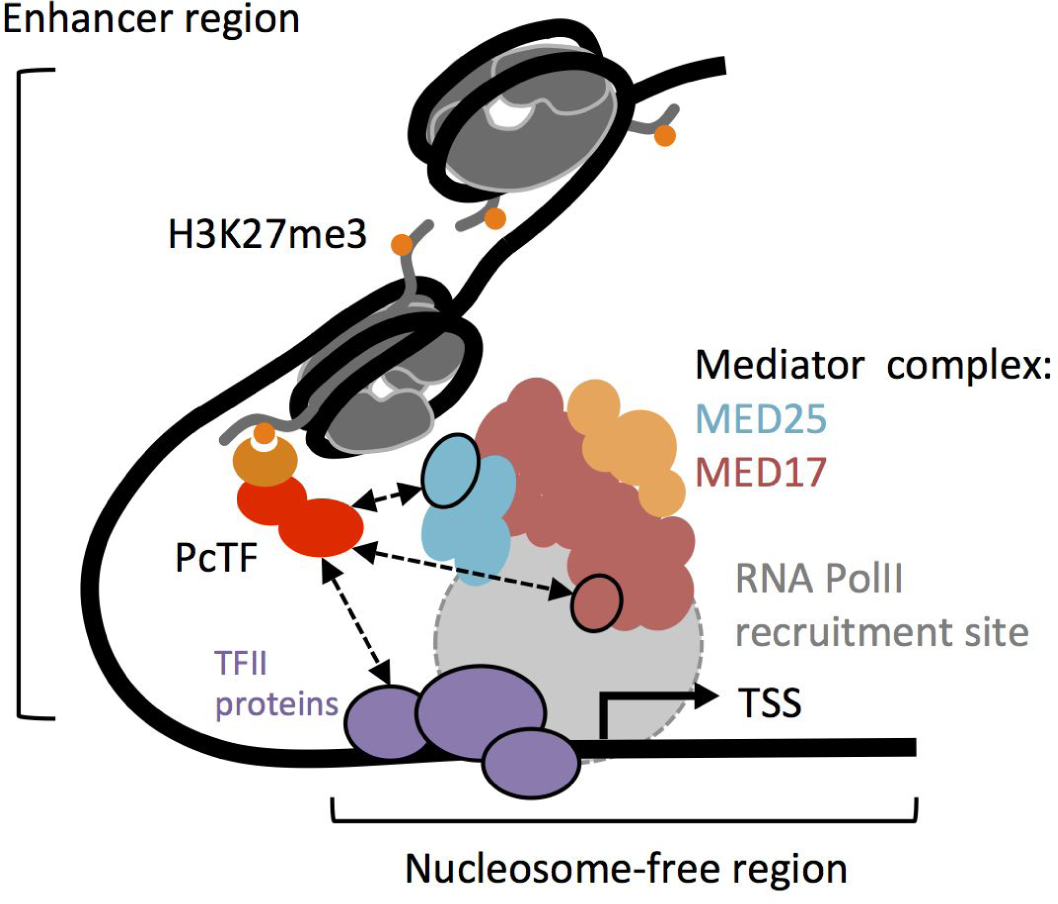
Model for PcTF-mediated gene activation. PcTF binds to H3K27me3 (orange dots) through the N-terminal PCD domain (orange crescent) at an enhancer region that is distal to the transcriptional start site (TSS). The C-terminal VP64 domain (red) interacts with members of the transcription initiation complex, including MED25, MED17, and TFII proteins. DNA is shown as a thick black line. The core histone complexes of nucleosomes and histone H3 tails are depicted in dark grey.

RNA-seq results for genes that do not appear to be direct targets of PcTF-mediated activation provide additional insights into gene regulation. Many of the genes we analyzed in the RNA-seq experiments do not change more than two-fold or become down-regulated after PcTF expression. In these cases, suboptimal positioning of the VP64 domain of PcTF relative to the promoter might neutralize its activity or lead to reduced expression below basal levels^31, 32^. We observed several genes that became up- or down-regulated, but lacked a PcTF signal within 10 kb of the TSS. Regulation of these genes could be mediated by transcription regulators that are direct targets of PcTF. For example up-regulated, PcTF-marked genes including *ATF3* and *CASZ1* encode broadly-acting transcriptional regulators that control multiple targets^43, 44^.

Our work has significant implications for medicine. Currently, epigenetic engineering relies largely on small-molecule inhibitors that are intended to erase cancer-associated epigenetic silencing and indirectly activate therapeutic genes. These drugs have major shortcomings such as incomplete erasure of cancer-associated methyl-histone marks and DNA damage. Here, we used a synthetic chromatin protein, PcTF, to convert the methyl-histone signal into direct activation of therapeutic genes instead of inhibitors to erase cancer-associated marks. Genes that were consistently upregulated by PcTF in all three cell lines we tested have therapeutic potential. Work by others has shown that induced upregulation of the transcription factor *ATF3* antagonizes cancer cell growth and invasion in certain tissues, such as ovarian (OVAC)^45^ and esophageal (ESCC)^46^. Overexpression of the *CASZ1* transcription regulator inhibits proliferation of neuroblastoma (SH-SY5Y)^47^. Additional work to determine the specificity of PcTF in cancer versus healthy cells and to enhance efficient synthetic protein delivery will bring this promising new technology closer to translation.

## ONLINE METHODS

### DNA constructs

The doxycycline-inducible PcTF-expressing plasmid has been described previously^14^. The plasmid is constitutively expressed in the absence of a TetR protein (in cell lines U-2 OS, SK-N-SH, and K562). The full annotated sequence of “hPCD-TF” is available online at Benchling - Hayneslab: Synthetic Chromatin Actuators (https://benchling.com/hayneslab/f/S0I0WLoRFK-synthetic-chromatin-actuators/seq-fSG2GUbr-hpcd-tf/).

### Chromatin Immunoprecipitation followed by deep sequencing (ChIP-seq)

Chromatin immunoprecipitation was performed on U-2 OS Flp-In T-Rex cells, which carry a chromosomal insert of a TetR-repressible PcTF gene as previously described^14^. A ChIP-Seq DNA Sample Prep Kit (Illumina IP-102-1001) was used to prepare deep sequencing libraries from DNA that was purified from immunoprecipitated (IP) and non-IP (input) chromatin. End-repair was carried out at 20°C for 30 min. in the following 50 µl reaction: 30.0 µl ChIP DNA, 1x T4 DNA ligase buffer, 0.4 mM dNTP, 1.0 µl T4 DNA Polymerase, 1.0 µl Klenow DNA polymerase, 1.0 µl T4 polynucleotide kinase. End-repair products were concentrated into a final volume of 34 µl Qiagen Elution Buffer (QIAquick PCR Purification kit 28104). 3’-end adenine base extension was carried out at 37°C for 30 min. in the following 50 µl reaction: 34.0 µl end-repair reaction product, 1x Klenow buffer, 0.2 mM dATP, 1.0 µl Klenow exonuclease. Base extension products were concentrated into a final volume of 10 µl H_2_O (Zymo Clean and Concentrator D4003). Adapter ligation was carried out at room temperature for 15 min. in the following 30 µl reaction: 10 µl 3’-base extension reaction product, 1x ligase buffer, 1.0 µl adapter oligo mix, 4.0 µl DNA ligase. Ligation products were concentrated into a final volume of 10 µl H_2_O (Zymo Clean and Concentrator D4003). 150 - 200 bp fragments from each ligation reaction were resolved and purified via gel electrophoresis and extraction (Zymoclean Gel DNA recovery kit D4001). DNA was back-eluted from the column twice with 10 µl H_2_O and brought to a final volume of 36 µl. Adapter-modified 150-200 bp fragments were enriched by the polymerase chain reaction (PCR) in the following 50 µl reaction: 36.0 µl gel-purified DNA, 1x Phusion buffer, 0.3 mM dNTP, 1.0 µl PCR primer 1.1, 1.0 µl PCR primer 2.1, 0.5 µl Phusion polymerase. The cycling program was 98°C/30 sec, 18x[98°C/10 sec, 65°C/30 sec, 72°C/30 sec], 72°C/5 min, 4°C/∞. PCR products were concentrated into a final volume of 15 µl H_2_O (Zymo Clean and Concentrator D4003). Libraries were size-confirmed on an Agilent 2100 Bioanalyzer and subjected to deep sequencing with single-end 100 bp reads on an Illumina Hi-Seq SR flow cell platform.

### Cell culture and transfection

Complete growth media contained 10% tetracycline-free fetal bovine serum and 1% penicillin and streptomycin (pen/strep). U-2 OS and U2OS-PcTF cells were cultured in McCoy’s 5A. K562, and SK-N-SH cells were cultured in IMDM, or EMEM, respectively. Cells were grown at 37 °C in a humidified CO_2_ incubator. U2OS-PcTF stable cell lines were generated by previously published work^14^. PcTF-expressing U-2 OS, K562, and SK-N-SH cells were generated by transfecting 5x10^5^ cells in 6-well plates with DNA/Lipofectamine complexes: 2 μg of plasmid DNA, 7.5 μl of Lipofectamine LTX (Invitrogen), 2.5 PLUS reagent, 570 µl OptiMEM. Transfected cells were grown in pen/strep-free growth medium for 18 hrs. The transfection medium was replaced with fresh, pen/strep-supplemented medium and cells were grown for up to 48 hrs. (K562) or up to 96 hrs. (U-2 OS and SK-N-SH).

### Quantitative Reverse Transcription PCR (qRT-PCR)

Total messenger RNA was extracted from ∼90% confluent cells (∼1-2x10^6^). Adherent cells (U2OS and SK-N-SH) were lysed directly in culture plates with 500 μl TRIzol. Suspended cells (K562) were collected into tubes, pelleted by centrifugation at 1000 xg for 3 min. at room temperature, separated from the supernatant, and lysed in 500 μl TRIzol. TRIzol cell lysates were extracted with 100 μl chloroform and centrifuged at 12,000 xg for 15 min. at 4 °C. RNA was column-purified from the aqueous phase (Qiagen RNeasy Mini kit 74104). SuperScript III (Invitrogen) was used to generate cDNA from 2.0 μg of RNA. Real-time quantitative PCR reactions (15 μl each) contained 1x LightCycler 480 Probes Master Mix (Roche), 2.25 pmol of primers (see Supplemental Table 1 for sequences), and 2 µl of a 1:10 cDNA dilution (1:1000 dilution for GAPDH and mCh). Crossing point (C_p_) values, the first peak of the second derivative of fluorescence over cycle number, were calculated by the Roche LightCycler 480 software. For each experiment (one cell type, one target gene, all time points) C_p_ values were normalized to the highest value. “Expression level” was calculated as delta C_p_, 2^[C_p_ GAPDH - C_p_ experimental gene], for PcTF-expressing cells. Fold-change was determined as Expression level divided by delta C_p_ for non-PcTF control cells.

### RNA-seq

RNA-seq was performed using two replicates per experimental condition. Total RNA was prepared as described for qRT-PCR. 50 ng of total RNA was used to prepare cDNA via single primer isothermal amplification using the Ovation RNA-Seq System (Nugen 7102-A01) and automated on the Apollo 324 liquid handler (Wafergen). cDNA was sheared to approximately 300 bp fragments using the Covaris M220 ultrasonicator. Libraries were generated using Kapa Biosystem’s library preparation kit (KK8201). In separate reactions, fragments from each replicate sample were end-repaired, A-tailed, and ligated to index and adapter fragments (Bioo, 520999). The adapter-ligated molecules were cleaned using AMPure beads (Agencourt Bioscience/Beckman Coulter, A63883), and amplified with Kapa’s HIFI enzyme. The library was analyzed on an Agilent Bioanalyzer, and quantified by qPCR (KAPA Library Quantification Kit, KK4835) before multiplex pooling and sequencing on a Hiseq 2000 platform (Illumina) at the ASU CLAS Genomics Core facility.

### Bioinformatics analysis

ChIP-seq alignments were carried out using the Bowtie2 algorithm^47^ with the hg19 reference genome (Feb. 2009 GRCh37). Enrichments normalized to input were calculated using the Hotspot algorithm (distribution version 4)^48^. ChIP-seq data for SK-N-SH was provided by B. Bernstein and K562 data was retrieved from the UCSC Genome Browser website^49^. Galaxy (http://www.usegalaxy.org) ^50^ was used to identify overlaps with a minimum of 1 bp between gene intervals and ChIP-seq enrichment intervals. ChIP-seq profiling across genes and intergenic regions was performed using the Cistrome platform^51^. RNA-seq alignments were carried out with de-multiplexed 50-bp single-end reads and the hg19 transcriptome using Bowtie 2. Transcript abundance and differential expression were calculated using Tophat and Cuffdiff^52^ within the Galaxy online platform using the *Homo sapiens* UCSC hg19 gene transfer format (gtf) file from Illumina iGenomes. Outdated genes were identified by cross-referencing gene symbols with the NCBI database. Out-dated genes and duplicate transcription start sites were removed from the Cuffdiff output in Excel, resulting in 23,245 genes. Analysis of distances between features (TSS’s, ChIP signals, and enhancers) was performed and all graphs and charts were generated in Excel.

## ACKNOWLEDGEMENTS

D. Nyer was supported by the Arizona Department of Health Services. D. Vargas was supported by the Western Alliance to Expand Student Opportunities (WAESO). C. Hom was supported by the Fulton Undergraduate Research Initiative (FURI). K.A. Haynes was supported by the NIH NCI (K01 CA188164) and the NSF Synthetic Biology Research Engineering Center (SynBERC). DNA oligos were purchased with support from Integrated DNA Technologies, a SynBERC IAB member. The authors thank J. Kramer, J. Steel, J. Park, and N. Briones from the Biodesign Institute at ASU for assistance with ChIP-seq and RNA-seq, and B. Bernstein for SK-N-SH ChIP-seq data.

